# Local auxin biosynthesis promotes shoot patterning and stem cell differentiation in Arabidopsis shoot apex

**DOI:** 10.1101/819342

**Authors:** Shalini Yadav, Harish Kumar, Ram Kishor Yadav

## Abstract

Shoot apical meristem (SAM) of higher plants is comprises of three distinct functional zones. Central zone (CZ) is placed at the meristem summit and harbors pluripotent stem cells. Stem cells undergoes cell division within the CZ and give rise descendants which enters into the surrounding peripheral zone (PZ), where they get recruited into organs. Stem cell daughters that get pushed underneath the CZ form rib meristem (RM). RM cells differentiate into stem tissue and vascular bundles. Understanding how stem cell daughters differentiate into PZ and RM cell types is essential to unravel the mechanism of meristem development in higher plants. Here, we show that meristem patterning and lateral organ primordia formation, besides intercellular transport, are also regulated by auxin biosynthesis mediated by two closely related genes belonging to *TRYPTOPHAN AMINOTRANSFERASE* family. In Arabidopsis SAM, *TAA1* and *TAR2* are required to maintain auxin responses and identity of PZ cell types. Furthermore, our genetic analysis shows that in the absence of local auxin production and transport differentiation of stem cells into PZ and RM cell types is stalled causing a complete arrest of shoot growth and development. Our study revealed that auxin biosynthesis and transport together control the patterning of SAM into PZ and RM cell types.

## Introduction

In Arabidopsis, cell-cell signaling network controlled by the receptor like kinase CLAVATA1 (CLV1) and its ligand CLAVATA3 (CLV3) regulates stem cell proliferation and exit in the shoot apical meristem (SAM) (Fletcher et al., 1999). CLV3 binds to CLV1 and initiates the signaling cascade (Ogawa et al., 2008), which restricts the expression of *WUSCHEL* (*WUS*) in the organizing center (Brand et al., 2000; Mayer et al., 1998). WUS protein moves from its site of synthesis to the CZ through plasmodesmata and directly activates *CLV3* and also represses genes involved in differentiation (Daum et al., 2014; Yadav et al., 2011; Yadav et al., 2013). Thus, WUS not only determine the fate of stem cells in the apical meristem but also serves as a central hub around which gene expression patterns are established to specify CZ, PZ, and RM to form a functional SAM.

Stem cell daughters are recruited into organ primordia at a regular interval in PZ. Past studies have shown that auxin, which is polarly transported by its efflux carrier PIN FORMED1 (PIN1), is required for this transition (Gälweiler et al., 1998; Reinhardt et al., 2003; Vernoux et al., 2000). By combining genetics, molecular biology, and pharmacological treatments, studies have shown that auxin controls its transport by regulating the expression of PIN1 and downstream auxin signaling network genes. Genetic and molecular evidence collected thus far suggests, PIN1 is polarized toward the regions of high auxin concentrations, reinforcing a positive feedback loop between PIN1 and auxin signaling network genes (Benková et al., 2003). A recent study has shown that auxin response factor *MONOPTEROUS* (*MP*), which controls the polarity of PIN1 non-cell autonomously, expresses in the incipient primordia first, PIN1 polarization follows MP expression (Bhatia et al., 2016). Although, *pin1* mutant plants produce cotyledons and leaves in vegetative phase of plant development (Vernoux et al., 2000), suggesting that locally produced auxin might be involved in leaf initiation in *pin1* mutant. Past models of phyllotactic pattern formation did not consider the role of locally produced auxin with transport in patterning of SAM.

Auxin is mainly synthesized from L-Tryptophan (L-Trp) in plants (Ljung, 2013). The enzyme encoded by *TRYPTOPHAN AMINOTRANSFERASE OF ARABIDOPSIS 1* (*TAA1*) and its close homologue *TRYPTOPHAN AMINOTRANSFERASE RELATED* gene (*TARs*) catalyse the conversion of L-Trp into indole-3-pyruvic acid (IPyA); subsequently, the YUCCA (YUC) monooxygenases convert IPyA to indole-3-acetic acid (IAA) (Zhao, 2010). Experimental evidence suggests that TAA1/TARs and YUCCA class of enzymes are the primary producer of IAA in Arabidopsis. The single mutant of TAA1/TARs and YUCs does not give phenotypes owing to their high genetic redundancy. However, when higher order mutants are generated, they display a defect in embryonic and postembryonic development related to auxin signaling (Cheng et al., 2007; Stepanova et al., 2008). Except for *TAA1*, it is not clear from present studies where and when the *TARs* genes are expressed in the SAM and whether their activity contributes in maintenance to local auxin maxima in the SAM is not explored.

Recent studies implicated the role of locally produced auxin in root meristem patterning. Expression studies revealed that *YUC1, YUC2, YUC4* and YUC6 are active in the SAM (Cheng et al., 2006). In contrast, expression of only *TAA1* is reported in few cells in the epidermal cell layer of SAM. These studies raise an important question, how do YUCs function in the absence of IPyA? This observation also raises the possibility of IPyA movement to YUCs expressing cells to achieve local auxin maxima.

Here, we report that *TAA1* and *TRYPTOPHAN AMINOTRANSFERASE RELATED 2* (*TAR2*) expresses in the SAM. By genetic analysis, we have uncovered the role of *TAA1* and *TAR2* in maintaining optimum auxin responses and PZ cell identity in SAM. The function of *TAA1* and *TAR2* is masked by auxin transport in SAM. Furthermore, when auxin transport mutant was combined with *taa1* and *tar2* mutant, it revealed that *TAA1* is required for stem growth whereas *TAR2* is required for lateral organ formation. In the absence of auxin transport and biosynthesis, genetic pathways involved in meristem patterning, organ initiation and their patterning get inhibited, and this results in complete arrest of SAM growth and development.

## Results

### Auxin produced by IPyA pathway affects PZ size in SAM

Past studies have shown that auxin produced via the IPyA pathway is critical for lateral organ patterning in plants (Brumos et al., 2018; Stepanova et al., 2008). From these studies it was not clear how IPyA contributes to plant growth and SAM maintenance in Arabidopsis. To understand the role of *TAA1* and *TAR2* in shoot and flower development, and how it affects SAM growth and development, we identified the T-DNA insertion in *TAA1* and *TAR2* from segregating *taa1*−/− *tar2*+/− plants. To uncover their role, *taa1-1*, *tar2-1* single and *taa1-1 tar2-1* (here after they will be referred to as *taa1*, *tar2* and *taa1 tar2*) double mutant plants were grown with wild type (WT) Col-0 as control for 4-weeks, and shoots were examined under confocal microscope after removing the older floral buds for shoot patterning and phyllotactic defects. Confocal imaging revealed no discernible impact of *taa1* and *tar2* on the patterning of shoot apex (Fig. 1A-C; Fig. S1E-G). *taa1* and *tar2* single mutant plants displayed normal inflorescence meristems (IM) architecture and height, and were indistinguishable from WT (Fig. S1A-C and I-K). Interestingly, the *taa1 tar2* double mutant plants grew extremely slow compared to respective single mutants and control and showed compact IM when grown under identical growth conditions (Fig. S1D and L). The *taa1 tar2* double mutant SAMs when observed from top appeared like oval in shape, and were small in size compared to respective single mutant and control (Fig. 1D; Fig. S1H and M). We also observed that *taa1 tar2* double mutant plant SAMs occasionally look like a pin shape, and lateral organ showed radial patterning (Fig. S2B). In addition, organ outgrowth was perturbed in double mutant compared to control, and flower primordium showed long pedicel with semi oval floral meristem (Fig. S2A, C, D). Taken together, these results show that *taa1 tar2* mediated auxin biosynthesis is critical for SAM and lateral organ growth and their proper patterning.

**Figure 1.**
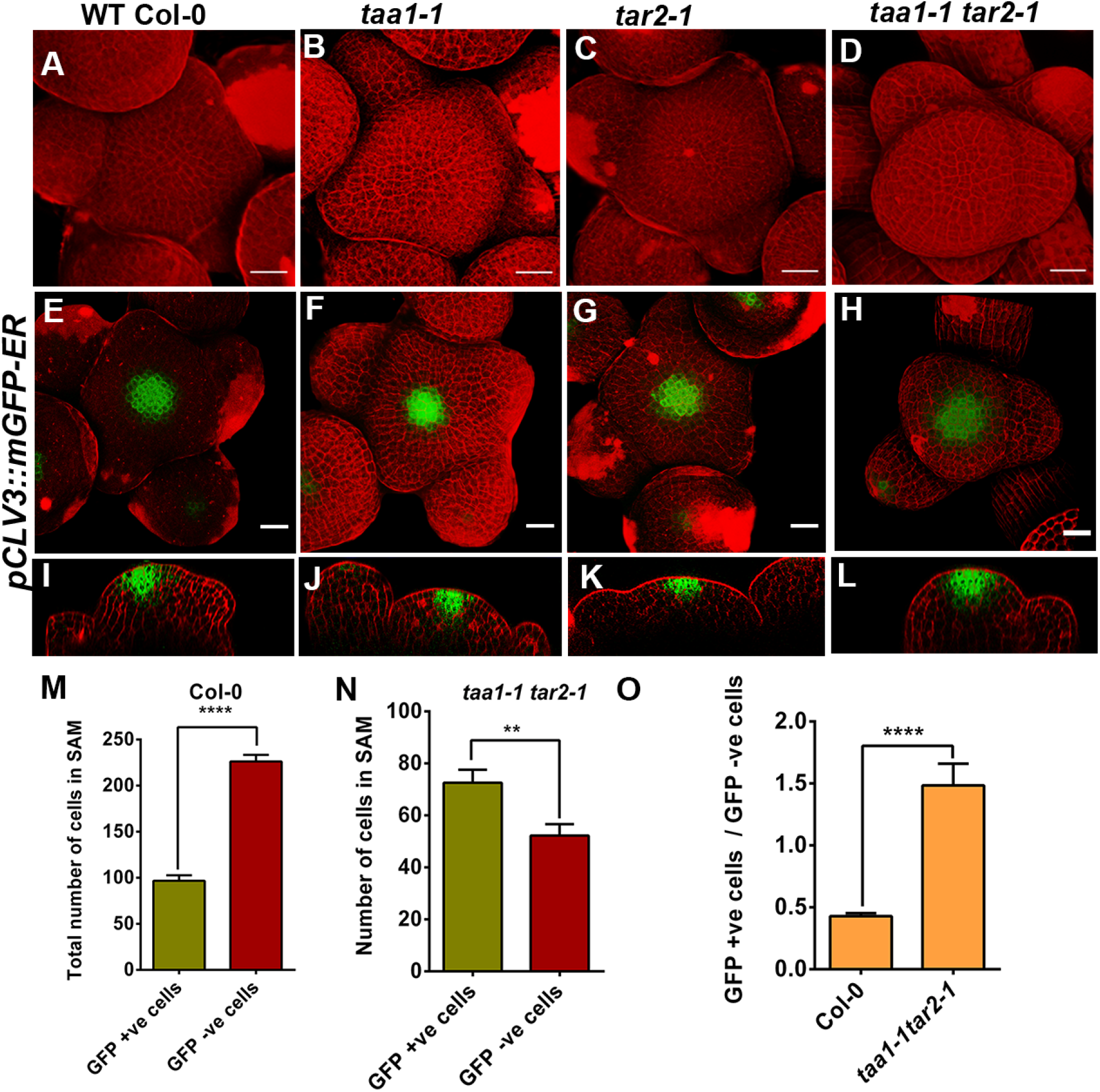
*TAA1* and *TAR2* are required for meristem development. Three dimensional (3D) reconstructed top view of L1 layer of SAM Col-0 (A), *taa1-1* (B), *tar2-1* (C) and *taa1-1 tar2-1* (D). Expression pattern of *pCLV3::mGFP-ER* in Col-0, *taa1-1, tar2-1* and *taa1-1 tar2-1*, respectively (E-L). 3D top view of SAM (E-H), and side view of the same image (I-L). Propidium iodide (PI, in red) was used to visualize cell outline. Quantification of *pCLV3::mGFP* +ve (CZ cells) and GFP −ve cells (PZ cells) in Col-0 (n=9) and *taa1-1 tar2-1* (n=9) double mutant (M-O), Error bars show SEM, asterisk marks represent statistically significant difference among the tested samples, Statistical test: Student t-Test (*p* < 0.005). Scale bars: 20μM. The raw data related to quantification of GFP +ve (CZ) and GFP −ve (PZ) cells as well as SAM size measurement are given in (Table S1). Scale bars = 20μM.

After uncovering that the *taa1 tar2* double mutant plants show slower growth and has smaller shoot size compared to control, we asked whether it is linked to reduced number of stem cells in them. Because a reduction in the rate of cell division and differentiation in CZ could affect the transition of stem cells into PZ cell types To test this, we made crosses between *taa1-1* −/− *tar2-1* −/+ with *pCLV3::mGFP-ER*, and isolated *taa1*, *tar2* and *taa1 tar2* double mutant plants carrying *pCLV3::mGFP-ER* reporter (Fig. 1E-L). A closer examination of *pCLV3::mGFP-ER* activity in *taa1 tar2* double mutant revealed that the number of cells showing GFP+ve signal did not reduce in single mutants and WT, suggesting that auxin produced by IPyA pathway do not affect CZ size. To evaluate the relative size of PZ, we counted both GFP+ve and GFP−ve cells in the epidermal cell layer of *taa1 tar2* double mutant as well as in single mutant and control SAMs (Table S1). This analysis revealed that *taa1 tar2* double mutant plants have smaller PZ compared to respective single mutant and control (Fig. 1M-O), indicating that auxin produced via IPyA pathway mainly affects PZ.

### Local auxin biosynthesis in meristem gives robustness to auxin signaling

Next, we asked whether the relative size of PZ in double mutant was affected due to a change in auxin signaling. To understand the role of local auxin biosynthesis in auxin responses in the SAM, we wanted to know whether the input and output responses of auxin are altered in the *taa1*, *tar2* single, and *taa1 tar2* double mutant plants. To map auxin responses, we crossed the *pPIN1::PIN1-GFP*; *pDR5rev::3XVENUS-N7* line with *taa1-1* −/− *tar2-1* +/− plants. After making the reporter homozygous in the *taa1*, *tar2*, and *taa1 tar2* double mutant background, respectively, we examined the meristem expressing *pDR5rev::3XVENUS-N7* and *pPIN::PIN1-GFP*. *DR5* responses in *taa1* and *tar2* single mutant did not change compared to WT (Fig. 2A-C and E-G). In contrast, in *taa1 tar2* double mutant auxin output was significantly reduced in the PZ of SAM (Fig. 2D, H, Y; Table S2). Polarization of PIN1-GFP was observed in *taa1, tar2* single mutant like WT (Fig. 2I-K and M-O). However, in *taa1 tar2* double mutant PIN1-GFP expression was weak, and the number of incipient primordia showing polarized PIN-GFP was reduced to one (Fig. 2L and P).

**Figure 2.**
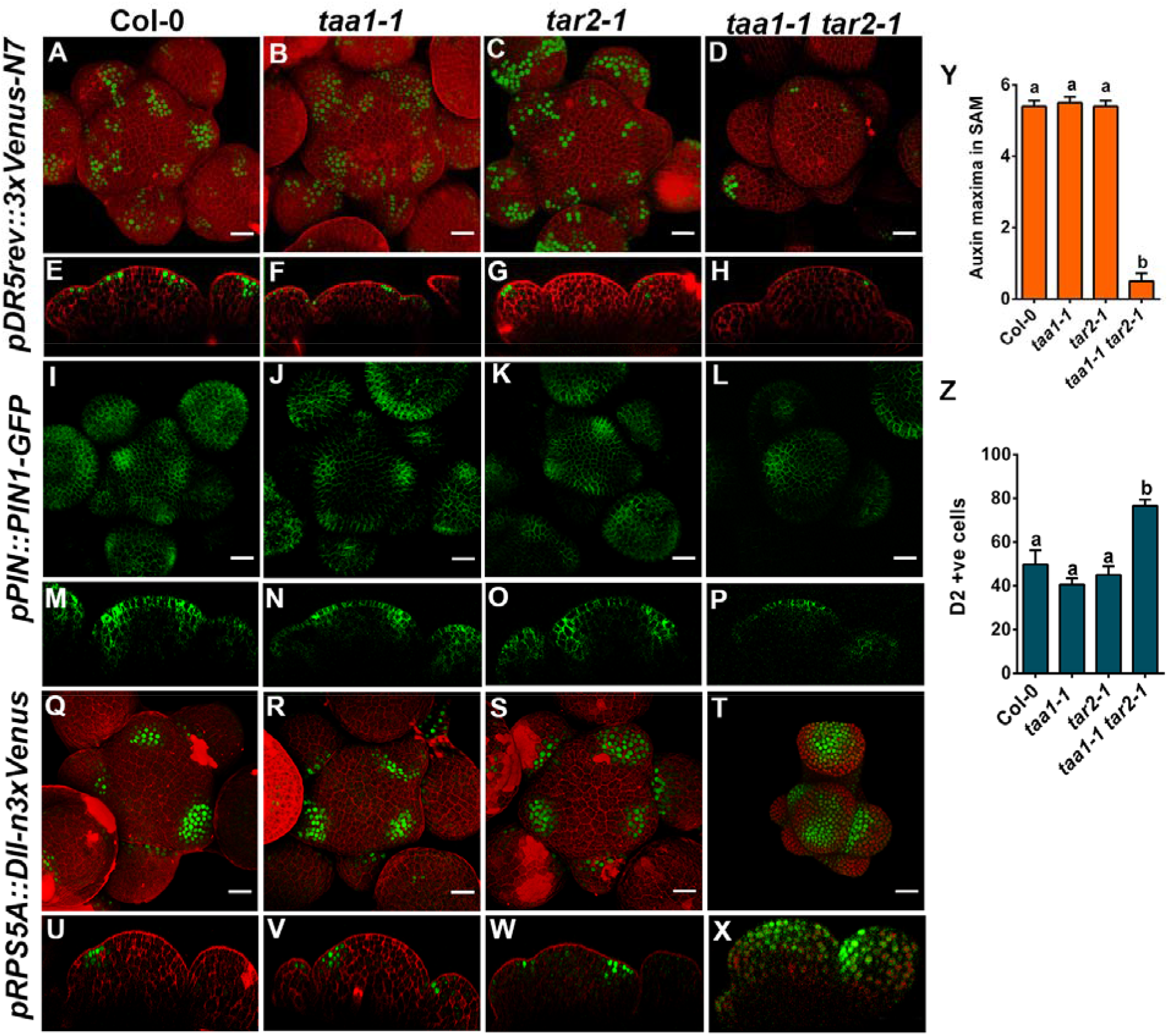
Auxin responses in SAM are dependent on auxin biosynthesis. 3D reconstructed top view of Arabidopsis SAMs showing the expression of the auxin output response *pDR5rev::3XVENUS-N7* in Col-0, *taa1-1, tar2-1* and *taa1-1 tar2-1* double mutant (A-D). Side view of (A-D) in (E-H). *taa1-1 tar2-1* double mutant SAM lacks *DR5* responses. SAM expressing *pPIN::PIN1-GFP* in Col-0, *taa1-1, tar2-1, taa1-1 tar2-1*, confocal stacks obtained from *pPIN1::PIN1-GFP* expressing plants were converted into 3D (I-P). Top of view SAM is shown (I-L), while the side of views of the same meristem is displayed in (M-P). Note, despite auxin biosynthesis defect, *taa1-1 tar2-1* double mutant plants show PIN1-GFP polarization and organogenesis. DII-Venus, auxin input sensor in Col-0, *taa1-1, tar2-1*, and *taa1-1 tar2-1* (Q-T). Top view of SAM L1 layer (Q-T), while the side views are displayed (U-X). Note, in *taa1-1 tar2-1* double mutant SAM, DII-Venus is stable despite weak auxin signalling. Quantification of auxin maxima in Col-0 (n=9), *taa1-1* (n=9) *tar2-1* (n=9) and *taa1-1 tar2-1* (n=9) (Y). Quantification of DII-Venus positive cells in Col-0 (n=8), *taa1-1* (n=8), *tar2-1* (n=8) and *taa1-1 tar2-1* (n=7) (Z). Statistical test: one-way ANOVA is followed by Tukey’s multiple comparisons test, different letters represent the statistically significant differences (*p* < 0.05). Quantification of data is given in (Table S2). Scale bars = 20μM.

Next, we checked the stability of auxin input sensor DII-Venus in the *taa1*, *tar2* single, and *taa1 tar2* double mutant to determine whether the local auxin biosynthesis mediated by *TAA1* and *TAR2* play any role in its stability in SAM. For this, we crossed the line carrying *R2D2* transgene with *taa1-2* −/− *tar2-1* +/−. For both single mutants as well as for double mutant plants were identified after following parents. The DII-Venus expression pattern was similar to WT in the single mutant of *taa1* and *tar2* (Fig. 2Q-S, U-W and Z). Interestingly, in the *taa1 tar2* double mutant plant SAM, DII-Venus protein appeared in the CZ cells of both IM and flower meristem, indicating the significance of *TAA1* and *TAR2* mediated auxin biosynthesis in maintaining an optimum threshold of auxin signaling not only in PZ but also in CZ of SAM (Table S2). Taken together, these results revealed that auxin biosynthesis mediated by IPyA pathway is essential for maintaining the optimum level of auxin signaling throughout the SAM. Despite weak PIN1-GFP expression, DII-Venus protein was stable in *taa1 tar2* double mutant SAM, indicating that auxin produced by *TAA1* and *TAR2* is essential in maintaining the optimum auxin input both in CZ and PZ cell types.

### *TAA1* and *TAR2* are expressed in the shoot and flower meristem

Our analysis of auxin output and input responses by *DR5* and DII-Venus in *taa1 tar2* double mutant showed that *TAA1* and *TAR2* are indispensable to maintain optimum threshold of auxin signaling in CZ and PZ cell types of SAM. Surprisingly, *taa1* and *tar2* single mutant plants do not show any phenotype. This could be due to their overlapping spatiotemporal expression pattern, and thus, they would complement each other. To understand the role of *TAA1* and *TAR2* in shoot and flower development, we examined the expression pattern of *TAA1* and *TAR2* by *in situ* hybridization in the inflorescence meristem (IM) of Arabidopsis. The *TAA1* transcript was detected in the epidermal cell layer of CZ cells in 4-week-old WT-L*er* SAM (Fig. 3A). In the sense probe, we did not see any signal (Fig. 3B). The expression pattern of *TAA1* was further investigated in IM using a translational fusion construct, where the *YPet* sequence was inserted upstream of the *TAA1* coding sequence (*pTAA1::YPet-TAA1*) to make an N-terminal fusion (Brumos et al., 2018). In comparison to the mRNA expression pattern reported by *in situ* hybridization, we found that the expression *pTAA1::YPet-TAA1* was slightly broad in the SAM but spatially restricted to the L1 layer (Fig. 3C). Similarly, we studied the mRNA expression pattern of *TAR2* by *in situ* hybridization in the 4-week-old shoot apices of WT-L*er*. In situ studies revealed expression of *TAR2* in the PZ of SAM, where the organs emerge in the shoot (Fig. 3D). Sense probe did not yield a signal (Fig. 3E). However, *TAR2* transcript conspicuously was missing from CZ and RM cells (Fig. 3D). To further investigate the expression of *TAR2*, we made a transcriptional fusion construct by amplifying the 3-kb promoter region of *TAR2* above the translational start site (TSS). We placed the 3-kb promoter fragment upstream to H2B-YFP translational fusion in a modified pGreen vector. Several independent T1 lines were selected to examine the *TAR2* expression in the shoot (n = 18). The majority of the reporter lines showed the activity of the *TAR2* promoter reminiscent to native mRNA expression pattern, as reported by in situ studies (Fig. 3F).

**Fig. 3.**
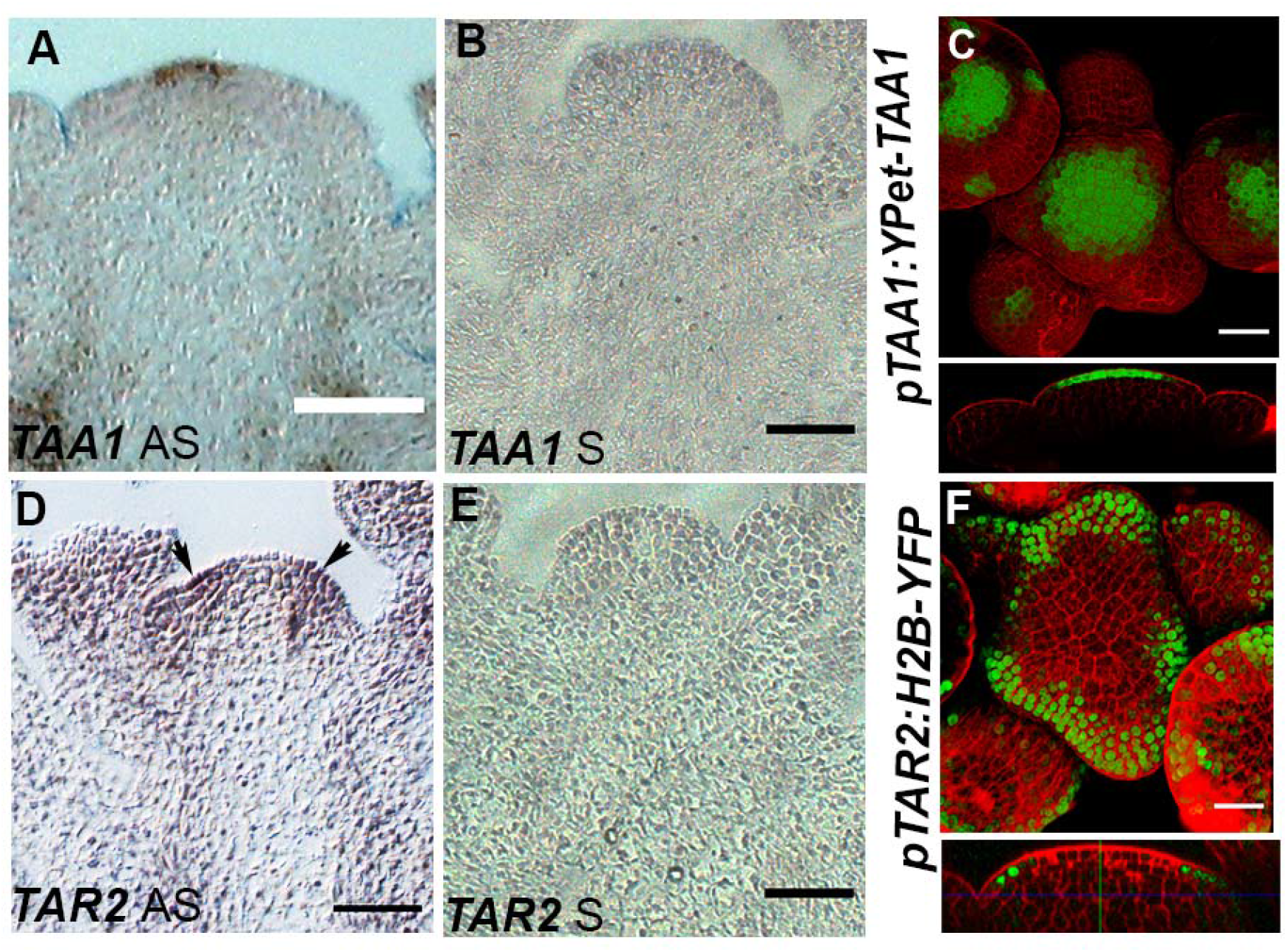
*TAA1* and *TAR2* are differentially expressed in SAM. *TAA1* expression is detected in the epidermal cell layer of SAM (A). *TAA1* sense probe was applied to the shoot apices in (B). Reconstructed surface projection of L1-layer of SAM expressing *pTAA1::YPet-TAA1* show an expansion in *pTAA1* activity along with side view (C). *TAR2* expression was recorded in the lateral organ primordia, as indicated by black arrow (D), in sense probe signal was not detected (E). Reconstructed of top and side views of SAM expressing *pTAR2::H2B-YFP* (F). Scale bars in A, B, D and E are 10μM, whereas in C and F, 20μM.

To understand when and where *TAA1* and *TAR2* mediated auxin biosynthesis is required in plant development, we checked the expression of *TAA1* and *TAR2* in embryo and seedling using the *pTAA1:: YPet-TAA1* and *pTAR2::H2B-YFP* lines, respectively. Both *pTAA1::Ypet-TAA1* and *pTAR2::H2B-*YFP did not show expression in the globular stage (16-32 cell stage) (Fig. S3A, J-K), however, at 64-cell stage Ypet-TAA1 fluorescence was visible in the epidermal cell layer of apical domain proembryo (Fig. S3B). Ypet-TAA1 is retained in the epidermal cell layer in the presumptive shoot apex of heart stage (Fig. S3C). In postembryonic stages, its expression was confined to the epidermal cell layer in SAM of 3-DAS seedlings and 4-week-old IM, respectively (Fig. S3-D-F). In stage-3 and 4 flowers, *TAA1* expression was observed in the center of the floral meristem (Fig. S3G-I). *TAR2* promoter activity was noticed first in the heart stage in the presumptive hypocotyl region of proembryo (Fig. S3L). In the postembryonic development, *TAR2* expression was seen in the periphery of SAM in 3-DAS seedlings (Fig. S3M, N). In flowers and IM, *TAR2* expression is extended beyond the L1 cell layer in the PZ cell types (Fig. S3O-R). Taken together, these results provide evidence that *TAA1* and *TAR2* show complementary expression patterns in embryonic and postembryonic development, and are regulated differentially in different stages of development. It is possible that IPyA produced by TAA1 and TAR2 can be transported to the neighboring cells where YUCs could catalyze its conversion into auxin, which would be transported more efficiently in and out of cells by diverse auxin transporter proteins. When the conversion of tryptophan into IPyA reduces significantly plants exhibits auxin related developmental defects.

### Auxin biosynthesis in the incipient organ primordia is critical for lateral organ initiation

Auxin biosynthesis defect in *taa1 tar2* double mutant plants affected the transition of stem cells into PZ cell types partially. We asked whether the lack of *DR5* activity or DII stability, if anyhow, is a good indicator of auxin signaling abrogation in SAM. We should see a complete repeal of lateral organ primordia formation in the *taa1 tar2* double mutant. We also reasoned based on the spatiotemporal expression pattern that the *TAR2* should be more critical in organ initiation than *TAA1*. To explore the role of PIN1 mediated auxin transport in organogenesis in conjunction with biosynthesis in SAM and how it affects the transition of stem cells into PZ cell types, we crossed the *taa1-1* −/− *tar2-1* +/− plants with weak *pin1-5* allele. Since strong alleles of *pin*, like *pin1-4* and *pin1-6*, do not form lateral organs in the reproductive phase despite having functional CZ, PZ, and RM. *pin1-5* mutant plants form lateral organs with defective flowers and set seeds (Fig. 4A, B, F, J; Fig. S4A, B). We isolated *pin1 taa1* and *pin1 tar2* double mutant plants, respectively, to ascertain the relative contribution of *TAA1* and *TAR2* based on their spatiotemporal expression pattern in auxin signaling and organogenesis (Fig. 4C, D; Fig. S4C, D). In the double mutant plants, the presence of the *pin1-5* allele was confirmed based on the flower organ number and flower morphology. To resolve the *taa1-1* and *tar2-1* genotype in the *pin1 taa1* and *pin1 tar2* double mutant, T-DNA PCR was setup for both *taa1* and *tar2*. The *pin1-5* single mutant inflorescences made at least 7-8 flowers (n=18), and flower primordia abut in the PZ similar to WT (Fig. 4F, J; and Table S4). Unlike strong alleles, the *pin1-5* allele also initiates and maintains axillary IM, which further bears the flowers and continues the growth (Fig. 4B, M, N; Table S3). In contrast, the main IM of both *pin1 taa1* and *pin1 tar2* double mutant plants bore a significantly lesser number of lateral organs before terminating in to pin like SAM (Fig. 4G, K, H, L, N-O; Table S5). *pin1 taa1*and *pin1 tar2* double mutant plants bear fewer flowers compared to *pin1-5* (Fig. 4N; Table S4), indicating that IPyA pool produced by TAA1 and TAR2 is critical for lateral organ formation in the SAM. The emergences of lateral organs further get reduced in *pin1 tar2* double mutant plant compared to *pin1 taa1*, suggesting that auxin produced in PZ has a more direct effect on the organ formation (Fig. 4D, N, O; Table S5). *pin1 taa1* double mutant plant axillary IM formed flower without developing pin-shaped apices at 35 days, indicating that the lateral organs are still being produced in the *pin1 taa1* double mutant while their *pin1 tar2* counterparts show exaggeration in pin-shaped apices formation (Fig. 4C, D). This genetic evidence makes *TAR2* more critical than *TAA1* in lateral organ formation in the reproductive phase of the plant development, which correlates well with its expression in the PZ of SAM. *TAA1* is required in the epidermal cell layer for apical dominance and stem tissue growth (Fig. 4C, D, P; Table S6). However, IPyA produced by TAA1 can partially rescues the lateral organ defect in the PZ in the absence of TAR2 (Fig. 4N, O; Fig. S4C).

**Figure 4.**
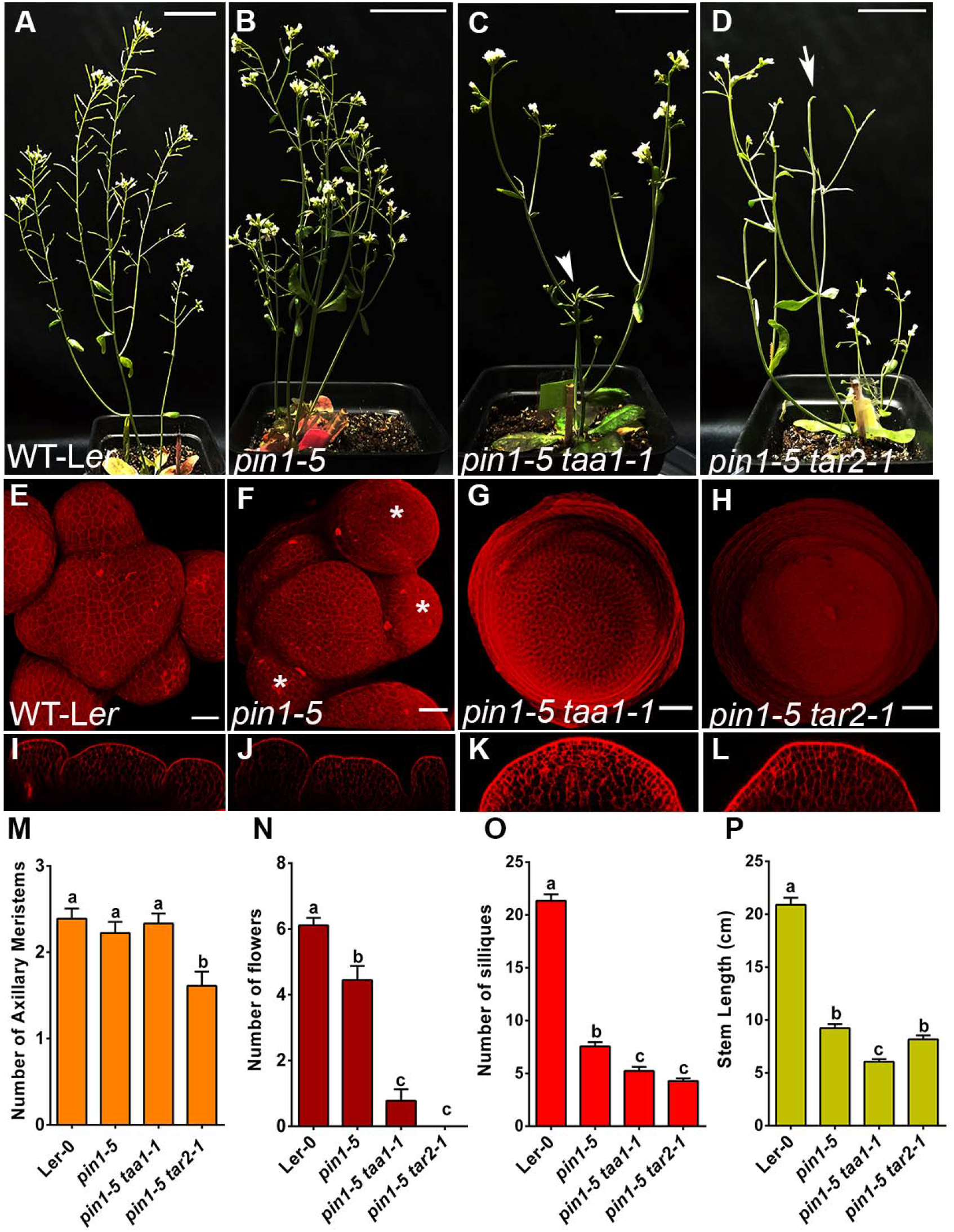
Auxin biosynthesis in emerging organ primordia is critical for organ initiation. From left to right 35-day old WT L*er*, *pin1-5, pin1-5 taa1-1*, and *pin1-5 tar2-1* (A-D). 3D reconstructed top view of WT L*er*, *pin1-5*, *pin1-5 taa1* and *pin1-5 tar2-1* SAM (E-H), and side views are given in (I-L). White arrowhead pointing towards the pin like inflorescence in *pin1-5 taa1-1*, and arrow in *pin1-5 tar2-1* plants. *pin1-5* makes lateral organs similar to WT (white asterisk). Graphs showing the quantification of axillary meristem, siliques numbers, flower numbers and stem length, respectively, WT L*er* (n=18), *pin1-5* (n=18), *pin1-5 taa1-1* (n=18) and *pin1-5 tar2-1* (n=18) (m-p). Statistical test: one-way ANOVA followed by Tukey’s multiple comparisons test, different letters represent the statistically significant differences (*p* < 0.005). Error bars show SEM. Scale bars: 20μM. quantification data for axillary meristem, flower numbers, silique numbers and stem height are given in supplementary (Table S3, S4, S5 and S6). Scale bars, A-D, 3cm, and for E-L, 20μM.

To understand the requirement of IPyA in rescuing the shoot phenotype in PZ of SAM, we replaced the H2B-YFP in *pTAR2::H2B-YFP* reporter construct with *TAR2* coding sequence. The resulting *pTAR2::TAR2* construct was transformed into *taa1*+/− *tar2* −/− genetic background, and independent lines rescuing *taa1 tar2* double mutant phenotype were identified by T-DNA PCR analysis (n=6) (Fig. S5A-F). The findings presented here suggest a role of auxin transport and biosynthesis in organogenesis and stem growth. Despite the redundant role played by *TAA1* and *TAR2*, their spatiotemporal expression pattern gives indications about their probable functions in stem tissue growth and lateral organ formation, respectively. Auxin transport mediated by PIN1 masks the effects of local auxin biosynthesis in *taa1* and *tar2* mutant background, suggesting that auxin transport and biosynthesis act in concert to promote SAM growth and development.

### Auxin signaling control the transition of stem cell progenitors into differentiating cells in PZ

To understand the role of auxin transport and biosynthesis on cell differentiation and phyllotaxy, we identified triple mutant plants for *pin1-5 taa1 tar2*. T-DNA PCR confirmed the *taa1* and *tar2* T-DNA alleles, which was verified by sequencing, while the *pin1-5* EMS mutant was sequenced directly to verify the mutation. The *pin1 taa1 tar2* triple mutant plants did not form rosettes in vegetative development (Fig. 5A-D). The majority of the triple mutant seedlings isolated from these crosses displayed unequal size cotyledons, and were rootless, but when kept for 2-3 week on soil in plant growth chamber, they formed a mound shape structure in between the cotyledons (Fig. 5A, C-D). Interestingly, triple mutant plants fail to produce leaves. We were unable to determine whether the stack of cells accumulated between the cotyledons was a mass of undifferentiated SAM tissue or regenerated stem tissue. To further investigate the role of auxin biosynthesis and transport in cell identity acquisition, we studied the expression of SAM marker genes by *in situ* hybridization in these plants.

**Figure 5.**
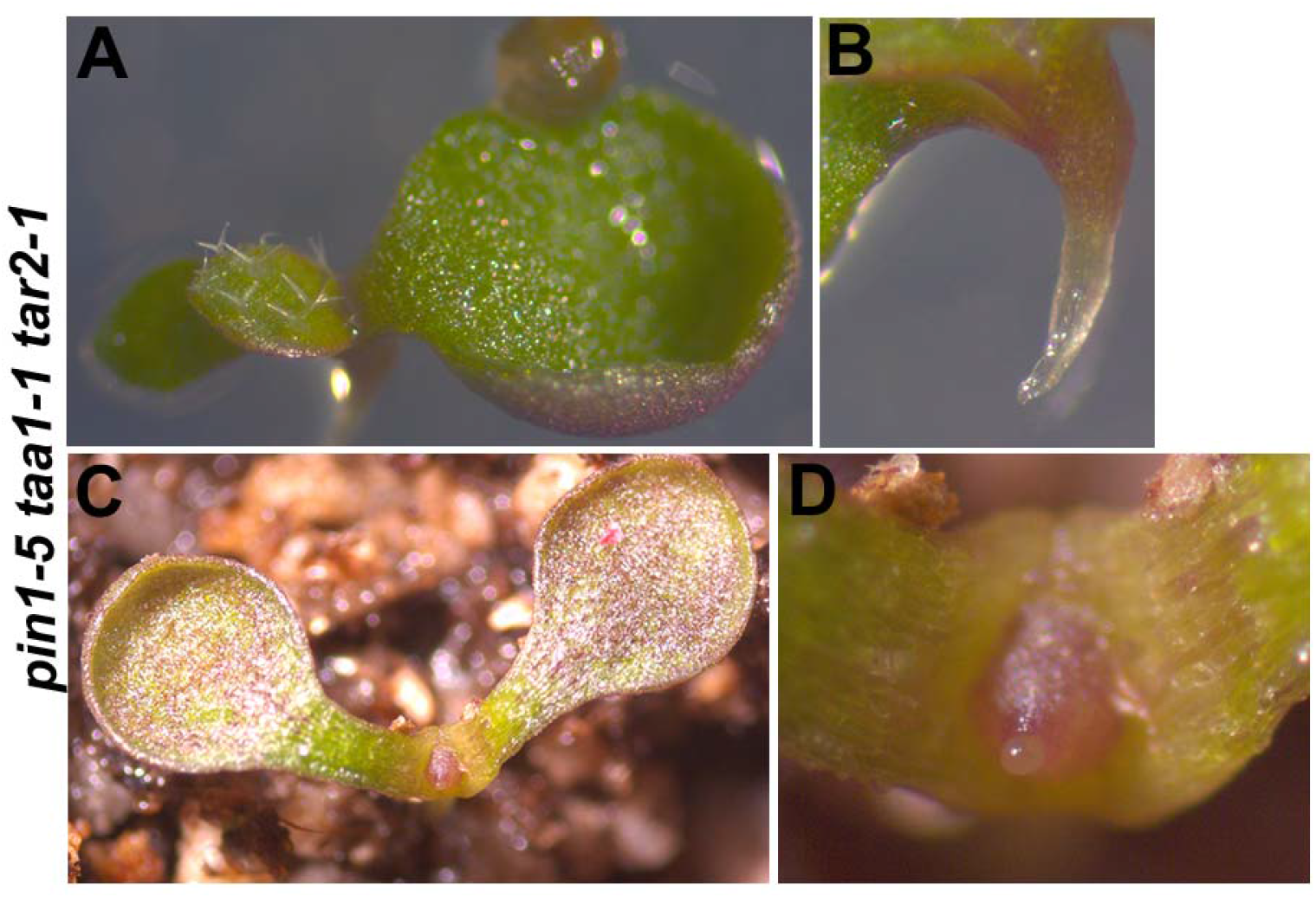
SAM growth and development is arrested when auxin biosynthesis mutant combined with transport. Triple mutant plants lacking functional PIN1, TAA1 and TAR2 show unequal size of cotyledons (A) and rootless seedling (B). Shoots emerge as an undifferentiated mass of cells without lateral organ and stem tissue in *pin1 taa1 tar2* triple mutant (C and D).

*CLV3* and *WUS* are usually expressed in CZ and RM cells, respectively (Fig. 6B, C). In *pin1 taa1 tar2* triple mutant plants, expression of *CLV3* mRNA was detected in the tip of the mound (Fig. 6Q). A closer examination of the in-situ images revealed a noticeable expansion of *CLV3* expression pattern in the epidermal and subepidermal cell layers towards the PZ in triple mutant compared to *taa1 tar2* double mutant (Fig. 6C, J). *WUS* expression level reduced dramatically in *pin1 taa1 tar2* triple mutant SAM compared to *taa1 tar2* double mutants, possibly due to overexpression of *CLV3* (Fig. 6I, P). To investigate whether these plants lack functional PZ or they fail to form the PZ cell type in the absence of auxin signaling, we examined the expression of marker genes, whose expression marks organ boundary regions and organ primordia in a functional SAM. The overall expression patterns of *ARF3*, *ARF4*, *ARF5*, and *CUC1* are maintained similar to WT in *taa1 tar2* double mutant (Fig. 6D-G, K-N). However, the level of expression has weakened significantly in *taa1 tar2* double mutant SAM (Fig. 6D-G, K-N). This has resulted in irregular organ boundaries and phyllotactic pattern formation and auxin signaling. In the *pin1 taa1 tar2* triple mutant, both the expression patterns and levels of *ARF3*, *ARF4*, *ARF5* and *CUC1* were either abolished entirely or reduced below the detection level, suggesting that auxin signaling is not only critical for timely transition of stem cells into differentiating cell types but also regulates the expression of essential CZ, PZ and RM genes (Fig. 6R-U).

**Figure 6.**
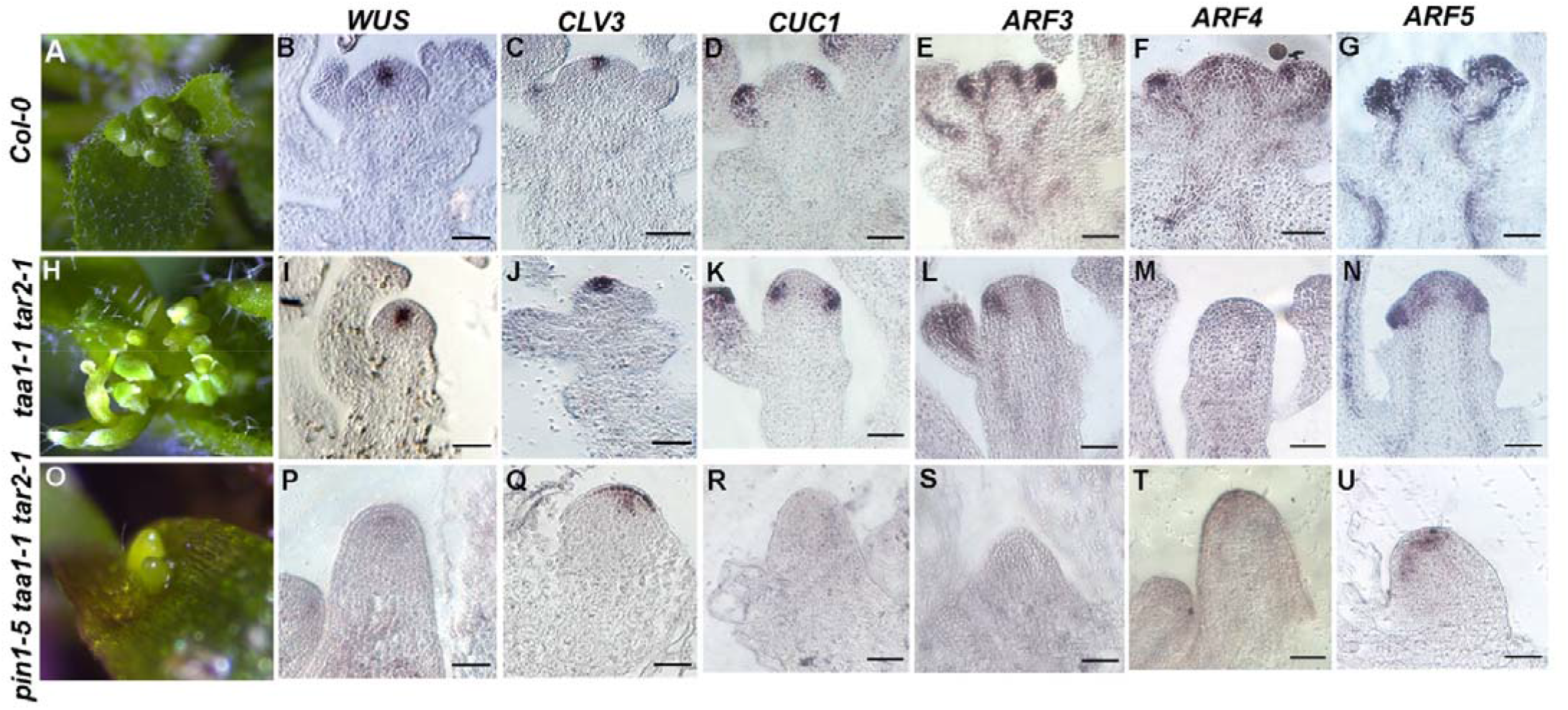
In the absence of auxin transport and biosynthesis patterning of SAM into PZ and RM cell types is stalled. Representative images of 4-week old shoot apex taken under dissecting scope for Col-0 (A), *taa1-1 tar2-1* double mutant (H), and two-week-old for *pin1-5 taa1-1 tar2-1* triple mutant plants (O). SAM marker genes for CZ (*CLV3*), PZ (*ARF3, 4, 5*), RM (*WUS*) and organ boundaries (*CUC1*) were analyzed in Col-0 (B-G), *taa1-1 tar2-1* (I-N) and *pin1-5 taa1-1 tar2-1* (P-U) by *in situ* hybridization. Note, expression of PZ and RM marker genes is either reduced or abolished in taa1 tar2 double and pin1 taa1 tar2 triple mutant. *CLV3* expression is expanded in towards the PZ in *pin1 taa1 tar2* triple mutant plants (Q). Scale bars = 10μM.

### Spatiotemporal regulation of auxin biosynthesis is required for shoot apex maintenance

We found that in the absence of auxin transport and biosynthesis functional organization of shoot apex was disrupted. And this has affected growth and development of SAM. Lateral organ initiation and their patterning were also compromised. Ideally optimum threshold of auxin signaling is maintained preferentially in the PZ because both biosynthesis and polar transport of auxin by PIN1 contributes in the auxin maxima formation. To test the influence of elevated levels of auxin on stem cell fate and meristem growth directly, we drove the expression of *TAR2* under the *35S* promoter. We generated *35S::TAR2* transgenic lines in WT L*er*. We found that the T1 plants did not show termination of the shoot apex (n=21) (Fig. 7A). However, when they were put again on soil in T2, eleven lines showed termination of the shoot apex in segregating plants (Fig. 7B, C). Semiquantitative RT-qPCR experiment revealed that plants showing shoot termination phenotype displayed elevated levels of *TAR2* transcript compared to WT control, indicating that beyond a threshold auxin levels in meristem can trigger cell differentiation (Fig. 7F). To test the impact of high auxin specifically on CZ and RM, respectively. We made a *6*×*OP::TAR2* operator construct and drove its expression using *pCLV3::LhG4* and *pWUS::LhG4* driver. Transformed selected for *pCLV3::LhG4; 6×OP-TAR2* showed termination of shoot after making 3-4 true leaves (n=48) (Fig. 7D). T1 plants carrying *pWUS::LhG4; 6×OP-TAR2* made only two true leaves (n=40) (Fig. 7E). Early onset of auxin biosynthesis in globular stage by *pWUS* correlates with the phenotype displayed by *pWUS::LhG4; 6×OP-TAR2* plants, while the *CLV3* promoter becomes active in heart stage. The delay in activation of *CLV3* thus allows plants to have 1-2 more leaves than *pWUS::LhG4; 6×OP-TAR2*. The data presented here shows that elevated level of auxin biosynthesis in SAM is detrimental for its maintenance. This also partly explains why *TAA1* and *TAR2* show complementary expression pattern.

**Figure 7.**
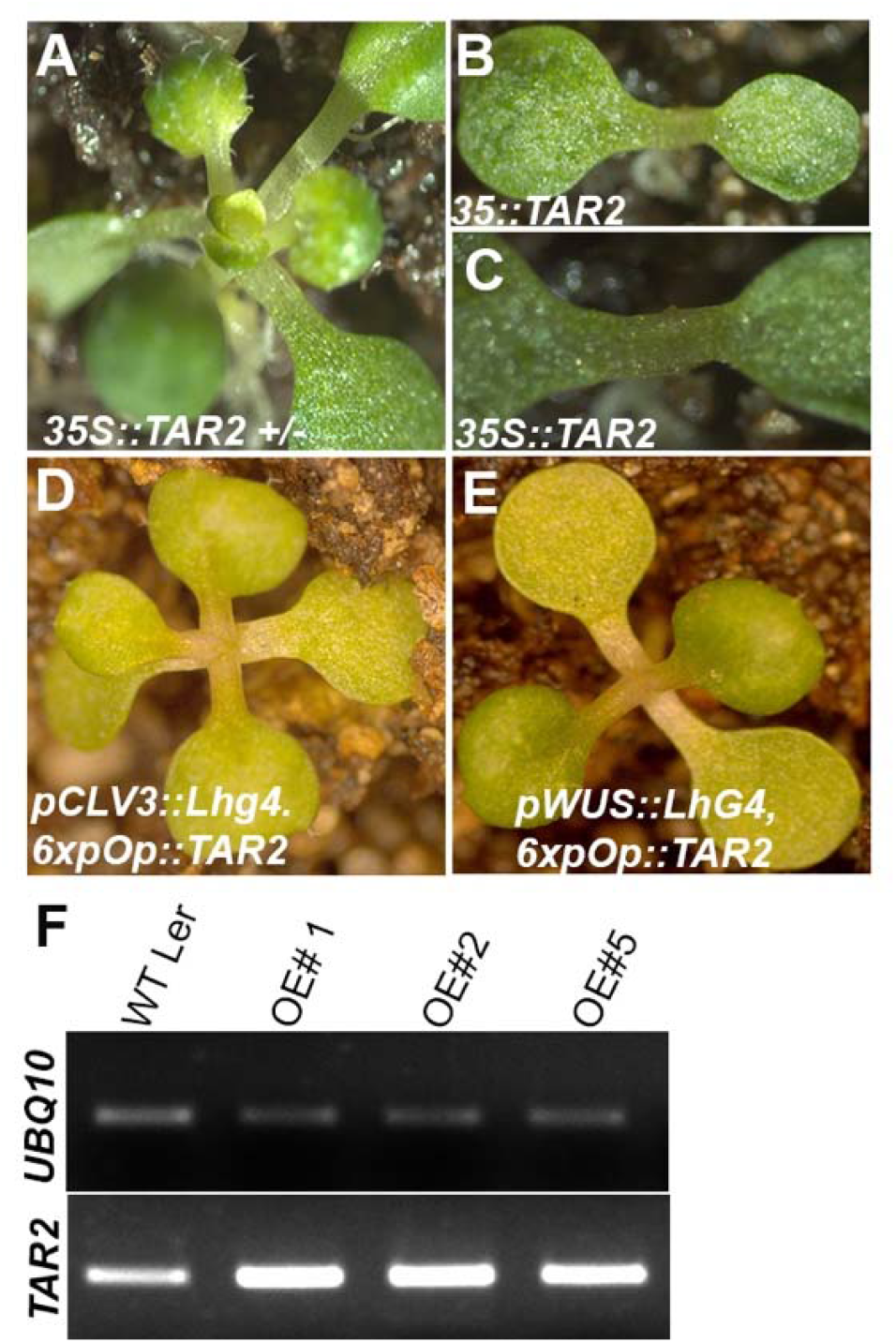
Ectopic expression of *TAR2* leads to termination of SAM. Twelve days old *35S::TAR2* +/− in T1 (A). In T2, they show termination of SAM (B and C). Ectopic overexpression of *TAR2* in *CLV3* and *WUS* domain cause termination of shoot (D and E). *TAR2* transcript in three independent lines quantified by semiquantitative qRT-PCR.

## Discussion

Despite of being a multifaceted phytohormone, auxin exquisitely regulates distinct aspects of plant development. In Arabidopsis shoot, cells of the developing organ primordia accumulate considerably higher amounts of auxin compared to neighboring cells that do not participate in primordium development. Considering the PIN1 mediated polar auxin transport, mathematical modeling studies explained this apparent disparity in auxin build-up and responses in the shoot apex. However, past studies were unable to explain why *pin1* mutant plants form lateral organs in the vegetative phase. Here, we show that disruption in the local auxin biosynthesis in SAM, mediated by *TAA1* and *TAR2*, leads to a decline in auxin signaling in SAM. This get weaken further when auxin transport is abolished in the biosynthesis mutants. The striking effect of this reduction in auxin signaling was on shoot patterning. *pin1 taa1 tar2* triple mutant plants failed to form organ primordia due to lack of PZ in these plants. In addition, stem cells also fail to differentiate into RM. Genetic evidence presented in this study strongly support the fundamental role of auxin signaling in SAM patterning.

*TAA1* and *TAR2* are upstream to YUCs in the IPyA pathways of auxin biosynthesis. In Arabidopsis SAM four YUCs genes are expressed. Despite their importance in achieving auxin maxima their overlapping expression patterns and high genetic redundancy makes it difficult to interpret the precise contribution of YUCs in auxin biosynthesis in distinct cell types of shoot. Past studies did not attempt to understand the role of *TAA1* and *TAR2* in achieving auxin maxima. IPyA is an important substrate for YUCs to produce auxin. In this study, we uncovered the role of *TAA1* and *TAR2* in stem growth and organogenesis. Our finding also confirms the role of auxin biosynthesis as reported by Cheng et. Al., (2006) and Cheng et. Al., (2007) in organogenesis including leaf development. In the absence of local auxin biosynthesis, optimum auxin responses in the PZ are abrogated. This is also attributed to the SAM patterning and phyllotactic defect observed in these plants.

Genetic evidence presented here show that organ boundary patterning is affected in the *taa1 tar2* double mutant and this genetic defect become more pronounced in *taa1 tar2 pin1* triple mutant, suggesting that auxin optima is not only critical for organ initiation but also required for organ patterning. Furthermore, *CUC1* expression is dependent upon the auxin. The expression of *CUC1* is progressively decreases in biosynthesis mutant when combined with transport mutant, resulting in complete arrest of organ formation. Our genetic data together with gene expression studies explains why *taa1 tar2* double mutant plants fail to pattern organ boundaries.

Studies have shown that auxin can activate auxin responsive genes by canonical binding to TIR/AFB co-receptor family proteins (Dharmasiri et al., 2005; Kepinski and Leyser, 2005), and noncanonical signaling mechanisms, directly binding to ETTIN/ARF3 transcription factor (Simonini et al., 2016). In both canonical and noncanonical signaling, auxin is the limiting factor. Of the six canonical auxin receptor protein encoded by the Arabidopsis genome, five are expressed uniformly in SAM, suggesting an intricate role in achieving the auxin signaling responses in the PZ cells (Prigge et al., 2020). Our data support a model where auxin emanating from PZ cells give positional information to stem cell descendants to differentiate into organ primordia. Previous studies have shown that expression of *ARF5* is dependent upon auxin signaling, and therefore, *ARF5* responds directly to canonical auxin signaling. However, these studies did not show how the expression *ARF3* and *ARF4* is regulated, and whether noncanonical signaling pathway is influenced by auxin levels. Our in situ studies on *taa1 tar2* and *taa1 tar2 pin1* shoot apices provide direct evidence that both locally produced auxin and transport controls the expression of *ARF3* and *ARF4* in SAM, thus auxin not only can regulate canonical signaling but also can regulate non-canonical signaling pathway mediated by *ARF3*.

## Materials and Methods

### Plant material and growth conditions

*Arabidopsis thaliana* Columbia-0 (Col-0) and Landsberg erecta (L*er)* ecotypes were used as WT strain and were obtained from Arabidopsis Biological Resource Center (ABRC, Ohio University, USA). T-DNA insertion alleles for *TAA1* and *TAR2* used in this study were obtained from ABRC. The *pin1*-5 allele is in L*er* ecotype and obtained from ABRC. R2D2 is in Col-0, and described in Liao et al., (2015), *pTAA1::TAA1-YPet* line is in Col-0, and was gifted by Anna Stepanova. *pRPS5A-DII-n3xVenus* is described by Brunoud et al., (2012). *pCLV3::mGFP-ER* line was available in house. *pPIN1::PIN1-GFP* and *pDR5rev::3xVenus-N7* are described in Heisler et al., (2005). Seeds were surface-sterilized with 70% ethanol, followed by 4% (v/v) sodium hypochlorite (MERCK 1.93607.1021) containing 0.02% Triton X-100 for 3-min and rinsed three times with sterile distilled water. The seeds were sown on Murashige and Skoog (MS) medium (Sigma) containing 0.8% Bacto agar (Himedia, India), 1% (w/v) sucrose and 0.1% (w/v) MES. Stratified seeds were kept in darkness at 4°C for three days and then transferred to plant growth chambers (Conviron, Canada, and Percival Scientific, USA). For transformation, WT L*er* and Col-0 dry seeds were sown on soilrite mix (KELTECH Energies Ltd. India) and kept for vernalization at 4°C for 3 days and then placed in the growth chambers (Conviron PGC Flex, Canada) under Philips fluorescent tube lights (120μmol light and 22°C) on a long day cycle (16 h light and 8 h dark). The soil was prepared by mixing soilrite mix (KELTECH Energies Ltd. India), compost, and perlite in the ratio of 3:1:1/2.

### Plasmid construct and generation of transgenic lines

To make *pTAR2::H2B-YFP* transcriptional fusion, *TAR2* promoter was PCR amplified using WT-L*er* genomic DNA as a template and cloned into pENTER/D/TOPO. A gateway LR-reaction was set up with a modified pGreen0229 destination vector (Yadav et al., 2010). *pTAR2::H2B-YFP* construct was introduced in the WT-L*er* to generate the reporter line. *35S::TAR2* construct was assembled in the pMDC32. For this, the *TAR2* coding sequence was PCR amplified and cloned into pENTER/D/TOPO. An LR reaction was set up between the pMDC32 destination vector and entry clone carrying the *TAR2* coding sequence. Resulting clones were confirmed by sequencing and transformed in WT L*er*. Transformants were selected on Hygromycin (50μg/ml). For ectopic expression of *TAR2* in CZ and niche cells. pENTER/D/TOPO vector carrying *TAR2* was used as an entry vector to setup LR reaction with *6×pOp* gateway compatible vector as reported in Yadav et al., (2010). Clones were sequenced and subsequently transformed into *pCLV3::LhG4* and *pWUS::LhG4* driver lines, respectively. Transgenics were selected on gentamycin (20μg/ml) and were transferred on the soil. Plants showed termination of the shoot apex in the vegetative phase and stayed green on soil for 2-3 week.

For rescue experiment, we made a new vector because of the *in planta* BASTA resistance of *taa1* T-DNA. To overcome this challenge, we assembled *pTAR2::TAR2* cassettes in pMDC32 vector. First, we replaced *35S* promoter with *pTAR2* inserts in *Sbf*I and *Asc*I restriction sites in pMDC32 vector. *Sbf*I and *Asc*I sites were introduced in *pTAR2* by PCR (Table S7). Secondly, pENTRY clone having *TAR2* CDS was used to setup the an LR reaction with *pTAR2* pMDC32. *pTAR2::TAR2* pMDC32 construct was transformed in *taa1-1* +/− *tar2-1*−/− and *pCLV3::mGFP-ER* reporter line by floral dip method. Seeds were collected and transgenics were selected on hygromycin (50μg/ml). More than 100 plants were selected on hygromycin and genotyped for *taa1-1* and *tar2-1* T-DNA insertion in T1 generation. In total 6 independent lines were identified for *taa1 tar2* double but looked similar to WT.

#### In situ Hybridization

Non-radioactive in-situ hybridization was performed according to the protocol posted on (http://www.its.caltech.edu/~plantlab/html/protocols.html). From the Arabidopsis WT cDNA library, coding regions of *CLV3, WUS, TAA1, TAR2, CUC1*, *ARF3, ARF4*, and *ARF5* were amplified and cloned into pENTER/D/TOPO vector (Table S7). Clones were sequence verified. To synthesize the full-length antisense and sense probe. First, the template was PCR amplified with the help of respective primer pairs (Table S7). One of the strands of the PCR product carries a T7 promoter sequence, which was used for in vitro sense and antisense probe synthesis, respectively (Yadav et al., 2010).

### Confocal Laser Scanning Microscopy of SAM

For confocal imaging, plants were first grown in 16 h light / 8 h dark conditions for four weeks. Once the bolting was induced, shoots were plucked with fine forceps and placed in 1.5% agar. Dissecting of the shoots was performed gently by removing the older flower buds using a stereomicroscope. To visualize the cell outline, shoot apices were stained with 10μg/ml propidium iodide (Invitrogen). The SAMs were scanned under the upright confocal microscope (Leica SP8, Germany). Images of 1024×1024 pixels were taken using a 63× long-distance water dipping lens with a step size of 1.5μm. The emission and excitation spectra were collected according to tagged fluorescence reporter. YFP, Venus, YPet were excited at 515 nm wavelength with argon laser lines at 9-10% laser power, and the emission spectra were collected between 524–540 nm by adjusting the variable band pass filter. GFP was excited at 488 nm, and emission spectra were collected at 500–530 nm. PI was excited using a 561 nm laser line, and emission spectra were filtered with a 600-650 nm adjustable bandpass filter. Transgenic lines established based on the inflorescence meristem expression pattern were used for embryo and seedling imaging. Siliques of appropriate age with immature seeds were dissected on a glass slide in distilled water for embryo imaging. Ovules were stripped from the ovary wall and placed into FM4-64 (Invitrogen) on a microscope slide. Embryos were popped out with the help of insulin syringes under a dissecting scope, and a coverslip was placed over the isolated embryos and sealed with nail polish. Imaging was carried out immediately under a 63× oil immersion objective in Leica SP8 upright confocal microscope as described above.

For seedling shoot apex imaging, seeds were surface sterilized with bleach and kept in the dark at 4°C for three days after putting on MS media plates. Plates were then kept vertically in the growth chamber for three days at 22°C. Germinated seedlings at their earliest stages (mostly with unopened/semi-opened cotyledons) were chosen for imaging and transferred on Magenta boxes containing 1.5% solidified agar. Seedlings were pushed into the pre-created holes into agar, leaving their top outside. SAM was exposed by gently removing one cotyledon with the help of fine tweezers (5TI, Dumont, Switzerland) and oriented vertically to visualize under the upright confocal nose piece. Propidium iodide (10μg/ml, Invitrogen) drops were put on the top of dissected seedlings and kept for 30-40 sec to visualize the cell outline. Autoclaved distilled water was poured over the seedlings to submerge, and z-stacks were taken by Leica SP8 upright confocal microscope equipped with a long-distance water dipping lens (63× objective). The rest of the confocal microscope settings were followed as described above.

#### Genetic analysis

T-DNA insertion mutants of *wei8-1 tar2-1* (CS16413) *tar2-1* (CS16404), and (CS69067) an EMS allele of *pin1-5* were obtained from Arabidopsis Biological Resource Center (ABRC, Ohio, USA). To confirm the insertion, first, genomic DNA was isolated using a modified CTAB method (Murray and Thompson, 1980). *taa1-1* single mutant was isolated from the segregating population of *taa1-1−/− tar2-1+/−*plants and genotyping was confirmed by T-DNA PCR, followed by sequencing using primers listed in Table S7. To generate the double mutants of *pin1-5 taa1-1, pin1-5 tar2-1*, and triple mutant of *pin1-5 taa1-1 tar2-1*, the single mutant of *pin1-5* was crossed with *taa1−/− tar2+/−* line. Seeds from the crosses were analyzed in F2 generation on the basis of genotype and phenotype. In F2, we identified *pin1-5 taa1-1, pin1-5 tar2-1, pin1-5 taa1-1 tar2-1*. Fecundity of *pin1-5 taa1-1* and *pin1-5 tar2-*1 double mutant plants was extremely poor despite that we were able to collect seeds and used them further to analyze the double mutant phenotype in subsequent generations.

To map the auxin input, output, as well as CZ size, *taa1-1−/− tar2-1+/−* mutant line was crossed with *pPIN1::PIN1:GFP, pDR5rev::3XVENUS-N7*, *R2D2*, and *pCLV3::mGFP-ER*, respectively. Plants from these crosses were propagated through F1. F2 seeds were collected separately for each independent line, and subsequently, 5 to 6 independent lines were followed in the F2 generation. Plants were genotyped for all the allelic combination *taa1-1−/−, tar2-1−/−, taa1-1+/− tar2+/−, taa1-1−/− tar2-1+/−, taa1-1+/− tar2-1+/−, taa1-1+/−tar2-1−/−, taa1-1+/−tar2-1+/+, taa1-1+/+ tar2-1+/−, taa1-1+/−tar2-1+/+, taa1-1-2+/+ tar2-1+/−, taa1-1+/+, tar2-1+/+* using the primers listed in Table S7. One independent line for each genotype was followed by F5 generation. Quantification of fluorescence +ve and −ve cell number for *pTAR2::H2B-YFP, pCLV3::mGFP-ER, pDR5rev::3XVENUS-N7*, and *R2D2* lines were performed using FIJI software.

### SAM size measurement

For SAM size measurements, confocal image stacks were obtained with 0.35μm step size from 4-week-old plants and analyzed using Morphograph X as described earlier by de Reuille et al., (2015). Original image in TIF format was loaded into stack1 of MorphographX (Fig. S6A). Gaussian Blur filter with 0.3 pixel radius was used to remove the noise (Fig. S6B). Signal edges were identified using edge detect process with a threshold value ranging from 10000 to 15000. Surface mess was created using Marching Cubes Surface with 2.5μm cube size (Fig. S6C). Visualized mess was subdivided two to three times (enough for extracting outer contour) after trimming the bottom. Mess was stretched 3-4 times with Smooth mess function with pass value of 20. Signal was projected on the mess with 2μm and 6μm top and bottom values, respectively. This resulting 2.5D image was used for the seeding for segmentation (Fig. S6C). One seed was used to mark the SAM boundaries excluding the emerging primordia by visual identification. Second seed was placed adjacent to the first seed, to restrict the propagation of first seed outside the meristem area (Fig. S6D). Watershed segmentation algorithm was used for the segmentation process (Fig. S6D). Since the segmented area was user defined and included only two propagated seeds, quality of segmentation was not an issue. Segmented area was converted into heatmap to get the actual values (Fig. S6E). Similar process was followed for all confocal stacks obtained for SAM measurements.

### Quantification of cells in SAM

Total number of *GFP* positive and *GFP* −ve cells were counted in the L1 layer from reconstructed top view of SAM as described by Yadav et al., (2010). Firstly, 3D stack of confocal image was loaded and opened into FIJI software directly and all the primordia were marked. Cells were counted by labelling manually in cell counter tool of FIJI software. Same protocol was followed up for quantification in mutant and WT SAM, respectively.

### Statistical analysis

All bar graphs were generated using GraphPad Prism 6 Software. Two tailed *t*-tests or one-way analyses of variance (ANOVA) followed by turkey’s multiple comparison test was done for statistical analysis using GraphPad Prism 6 software.

## Author Contributions

R.Y. conceived the project, S.Y. and R.Y. designed experiments. S.Y., H.K., and R.Y. performed the experiments. S.Y., H.K., and R.Y. analysed the data. S.Y. and R.Y. wrote the paper.

## Acknowledgment

We thank Anna Stepanova for *pTAA1::YPet-TAA1* seed lines, Teva Vernoux for *pRPS5A::DII-n3xVenus*, Dolf Weijers for R2D2 line, Thomas Laux for *pCLV3::LhG4* and *pWUS::LhG4* driver lines seeds. We thank Venu Reddy for sharing *pCLV3::mGFP-ER* lines. Imaging work was carried out at the confocal microscopy core facility of IISER Mohali. The IISER Mohali core grant funds support confocal imaging facility.

