## Supplement Information for "Local auxin biosynthesis promotes shoot patterning and stem cell differentiation in Arabidopsis shoot apex"

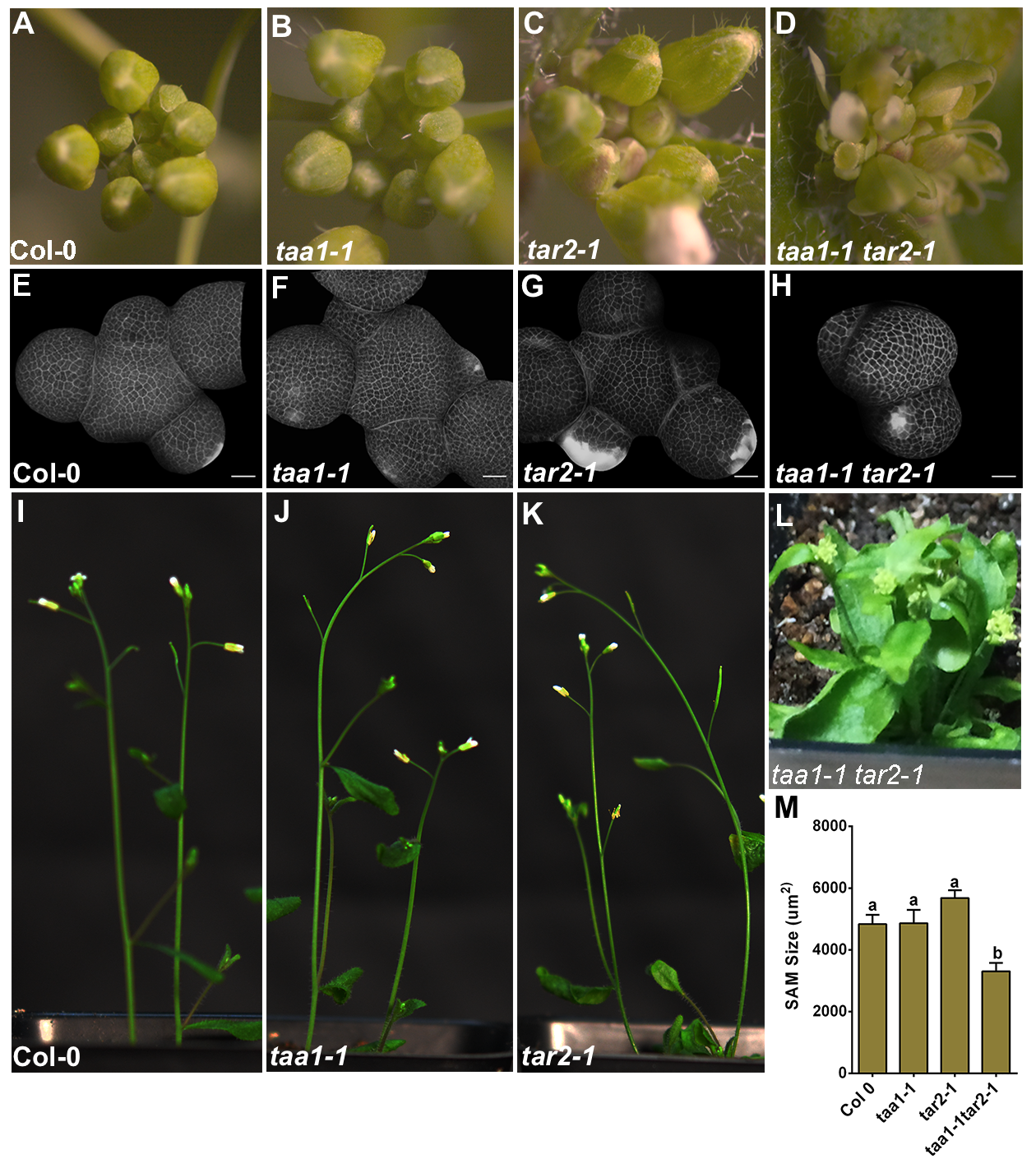
**Supplementary Figures**

**Fig. S1. *taa1 tar2* double mutant plant shows smaller shoot size.**

Inflorescence meristems (IMs) of 4 week old Col-0 (A), *taa1-1* (B), *tar2-*1 (C), and *taa1-1 tar2-1* double mutant (D). Three dimensional (3D) image of 4 weeks old Col-0 (E) *taa1-1* (F) *tar2-1* (G) and *taa1-1 tar2-1* (H) SAM. The double mutant shows growth defects and reduced SAM size. Plant height of four week old Col-0 (I) *taa1-1* (J) *tar2-1* (K) and *taa1-1 tar2-1* (L) was captured using camera. SAM size measurement carried out from confocal image stacks obtained from Col-0 (n=6), *taa1-1* (n=7), *tar2-1* (n=7) and *taa1-1 tar2-1* (n=7) plants (M, Table S1). Error bars show SEM. Statistical test: one way ANOVA followed by Tukey's multiple comparisons test, different letters represent the statistically significant differences (*p*<0.005).
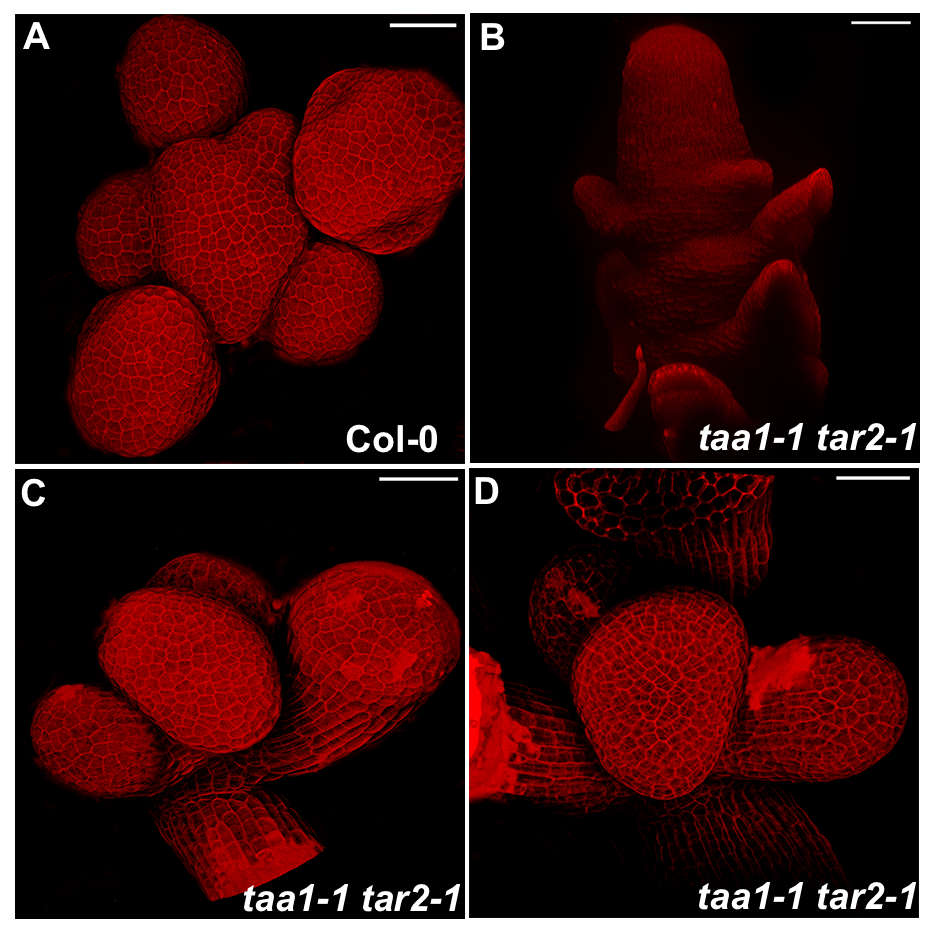
Scale bars; 20µm.

**Fig. S2**. ***taa1 tar2* double mutant plants show defect in organ patterning.**

Confocal image stacks were rendered in to 3D top view of SAM for Col-0 (A) and *taa1 tar2* double mutant SAMs (B-D). Scale bars = 20µM.

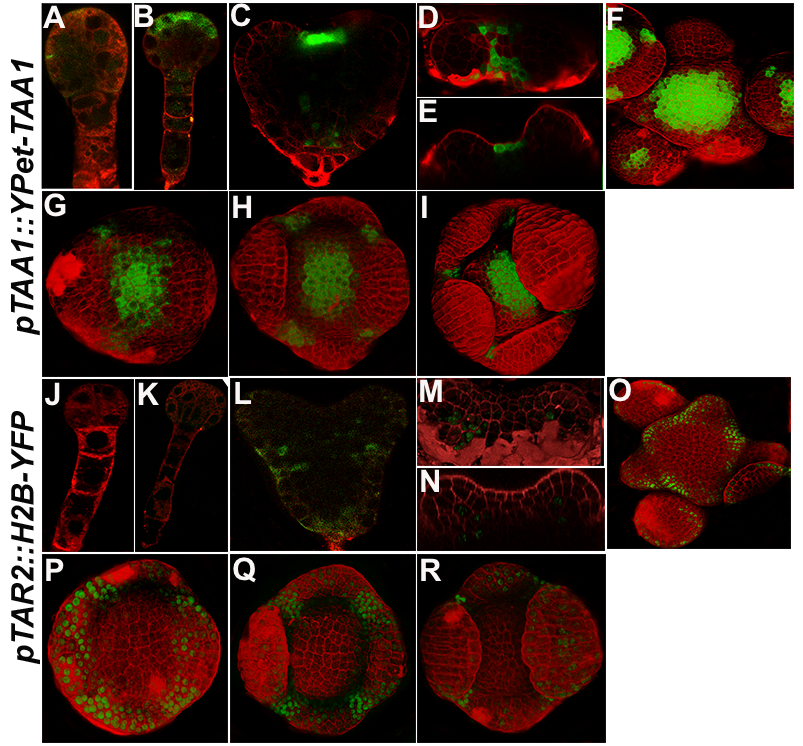

**Fig. S3. Expression pattern of *TAA1* and *TAR2* in embryo, seedling, and flower.** Confocal images of *pTAA1::Ypet-TAA1* expression in early globular (A), late globular (B), and heart stage (C) (in green). *TAA1* expression in 3 DAG seedlings, a top view is shown in (D) while side view is shown in (E). *TAA1* expression profile during different stages of flower development. *pTAA1::Ypet-TAA1* expression pattern observed in stage 3 and 4 flowers (F-I). *TAA1* expression is restricted to the epidermal cell layer in embryo, seedling, and IM. *TAR2* expression in the early globular stage (J), late globular stage (K), and heart stage (L). *TAR2* expression in 3 days old seedling restricted to PZ. Top view of 3-day old of SAM expressing *pTAR2::H2B-YFP* in (D) and side view in (E). *TAR2* expression is lacking in the floral meristem while emerging sepal primordia show a uniform expression pattern (P-R, stage 3, and 4 flower primordia). Cell outlines were stained with PI (in red). *TAR2* expression is restricted in the PZ in SAM from early seedling stage to IM.

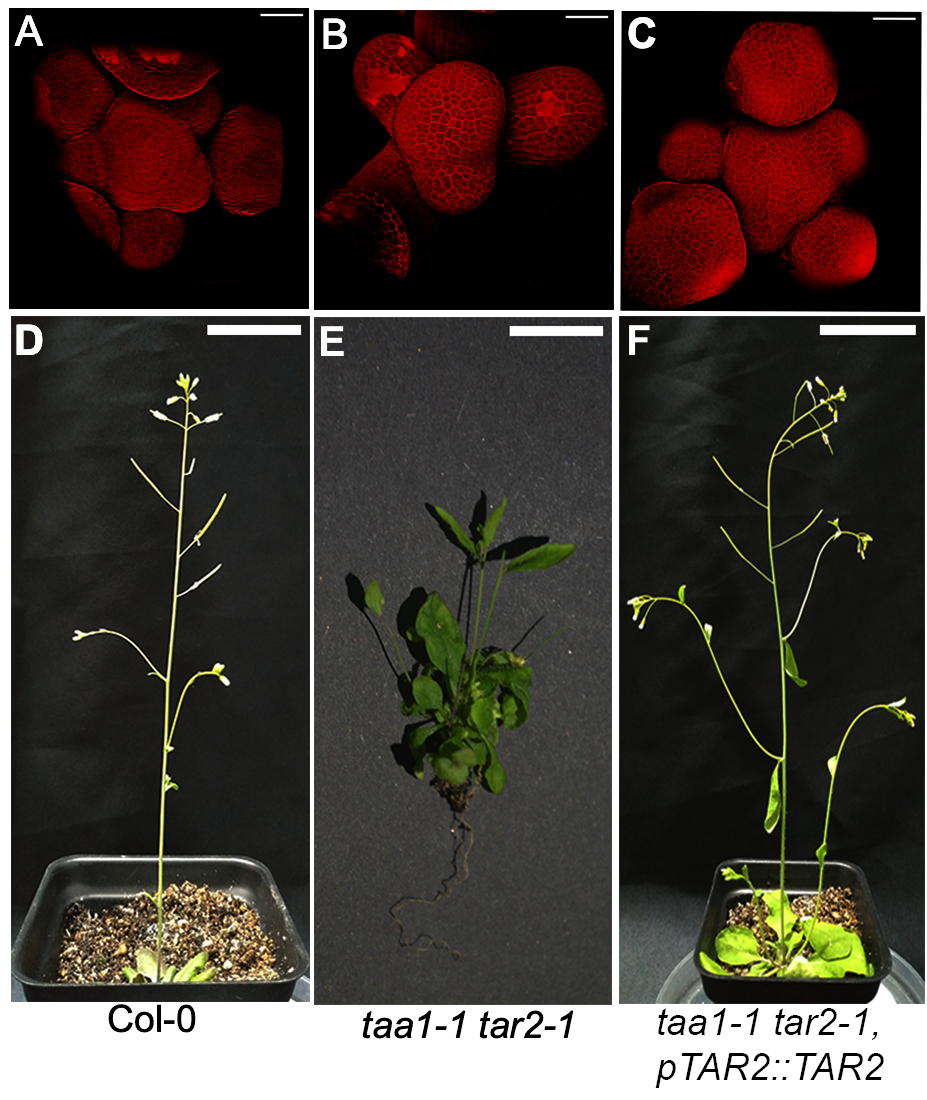

**Fig. S4. Defect in auxin transport and biosynthesis affect organogenesis.**

Images of twenty-eight-day old WT (A), *pin1-5* (B)*, pin1-5 taa1-1* (C) and *pin1-5 tar2-1* (D) plants were captured with camera*.*  *pin1-5-/- tar2-1-/- taa1-1+/-* plants terminate into pin-like SAM. Scale bars in A-C; 20µM and D-F, 3cm.

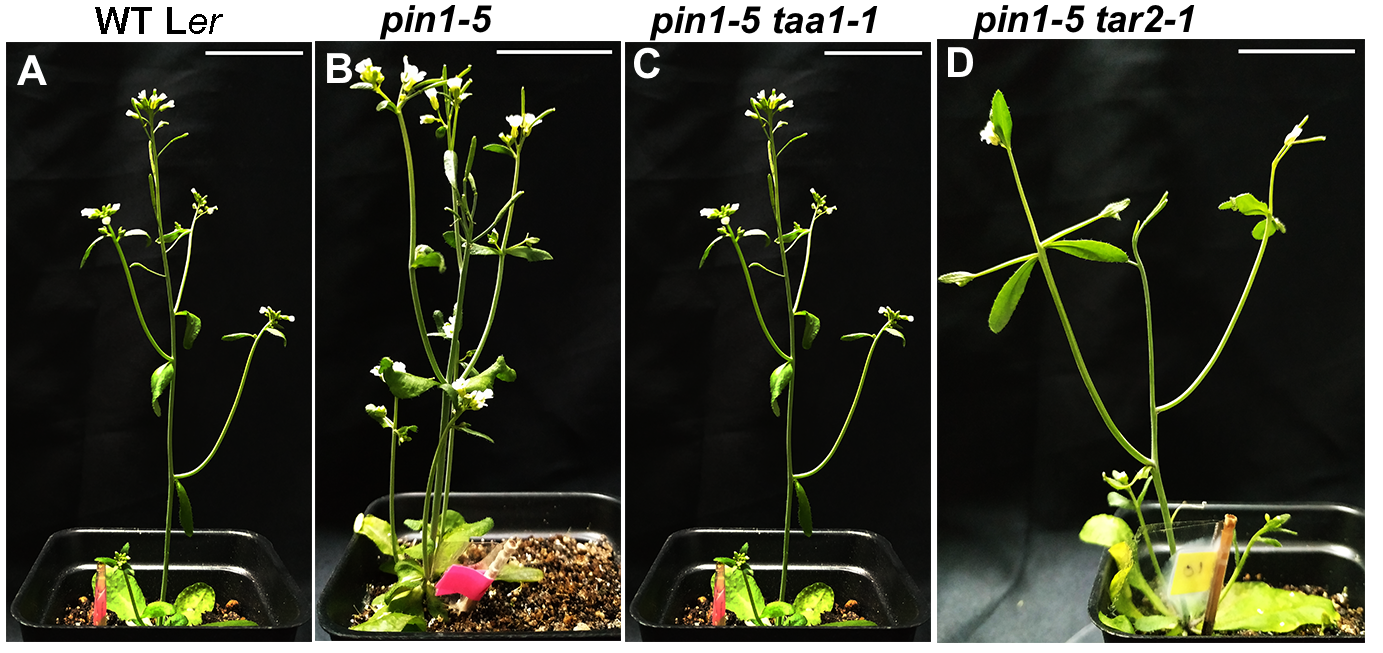

**Fig. S5. *pTAR2::TAR2* transgene rescues the *taa1 tar2* double mutant phenotype.**

3D reconstructed top view of 4-week old soil grown Col-0 (A), *taa1 tar2* double mutant (B), and *taa1 tar2* double mutant plant rescued by *pTAR2::TAR2* construct. Phenotypes of adult plants grown parallelly on soil WT Col-0 (D), *taa1 tar2* (E), and *taa1 tar2* rescued line. *pTAR2::TAR2* transgenes rescues the double mutant phenotype. Scale bar: 3cm

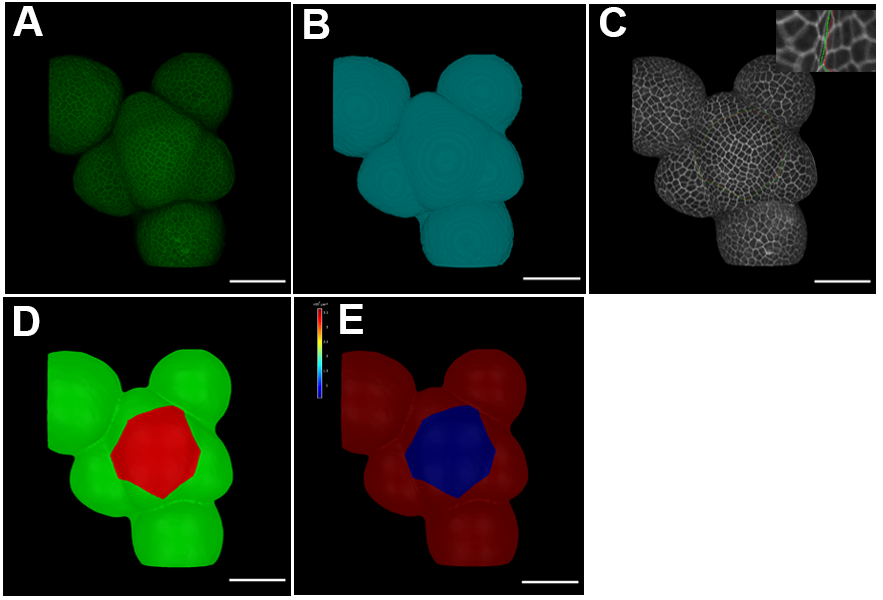

**Fig. S6. Methodology used for SAM measurement.**

Original image loaded into Morphograph X environment (A), surface detection after filtering the image (B), 2.5D reconstruction created by projecting the signal onto mess (C), Seed labelling to separate the organ primordia and SAM boundaries (D), Segmented image after watershed segmentation algorithm (E) and surface area Heat map obtained from segmented image (F). Scale bars = 50µM.
